# Integrating Gene Synthesis and Microfluidic Protein Analysis for Rapid Protein Engineering

**DOI:** 10.1101/025239

**Authors:** Matthew C. Blackburn, Ekaterina Petrova, Bruno E. Correia, Sebastian J. Maerkl

**Affiliations:** Institute of Bioengineering, School of Engineering, École Polytechnique Fédérale de Lausanne, Lausanne, Switzerland

## Abstract

The capability to rapidly design proteins with novel functions will have a significant impact on medicine, biotechnology, and synthetic biology. Synthetic genes are becoming a commodity, but integrated approaches have yet to be developed that take full advantage of gene synthesis. We developed a solid-phase gene synthesis method based on asymmetric primer extension (APE) and coupled this process directly to high-throughput, on-chip protein expression, purification, and characterization (mechanically induced trapping of molecular interactions, MITOMI). By completely circumventing molecular cloning and cell-based steps, APE-MITOMI reduces the time between protein design and quantitative characterization to 3-4 days. With APE-MITOMI we synthesized and characterized over 440 zinc-finger (ZF) transcription factors (TF), showing that although ZF TFs can be readily engineered to recognize a particular DNA sequence, engineering the precise binding energy landscape remains challenging. We also found that it is possible to engineer ZF – DNA affinity precisely and independently of sequence specificity and that *in silico* modeling can explain some of the observed affinity differences. APE-MITOMI is a generic approach that should facilitate fundamental studies in protein biophysics, and protein design/engineering.

---

Engineering proteins with novel functions remains a challenging task. Experimental methods that rely on generating large numbers of random protein variants followed by screening are currently the most successful approaches to protein engineering ^1^. Rational design of protein function, on the other hand, remains a major unsolved problem in biochemistry. Nonetheless, computational approaches are becoming adept at informing protein design and predicting function^2,3^, and techniques that permit the rapid generation and quantitative characterization of designed protein variants would greatly aid protein engineering and further improve the accuracy of computational approaches. Numerous gene synthesis methods have been developed^4,5^ and synthetic genes are becoming a commodity, but little to no attention has been given to streamlining the downstream processing steps required to quantitatively characterize the large number of proteins that can potentially be generated using gene synthesis based approaches.

Here we present a pipeline for the rapid synthesis and characterization of rationally designed synthetic proteins. We developed a bench-top solid-phase gene synthesis method based on APE, and demonstrate that expression ready linear templates generated by APE can be used directly for on-chip high-throughput protein expression, purification, and quantitative characterization by MITOMI^6,7^. APE synthesizes genes with fidelity comparable to the best currently available methods and can synthesize genes with internal sequence redundancies. APE-MITOMI completely circumvents any requirement for molecular cloning and cell-based protein expression, and can synthesize and characterize hundreds of novel protein variants per week. As a proof-of-concept we applied APE-MITOMI to the engineering and characterization of Cys_2_His_2_ ZF TFs. We found that ZF TF affinity can be precisely tuned independently of specificity and although it is possible to engineer specificity, the precise binding energy landscape is more difficult to rationally engineer.

Individual ZF domains provide a convenient structure for refactoring due to their relatively small size and composability^8,9^ (Fig. 1a). ZFs fused to nucleases (ZFN) are one of the main tools currently used for clinical genome editing^10,11^. The versatility of ZFs can be further expanded by fusing them to other effector domains allowing them to perform a variety of site-specific genetic modifications beyond DNA cleavage^12,13^. Synthetic ZFs have also been used to construct artificial transcriptional regulatory circuits in yeast^14,15^. ZF TFs are thus ideal targets for exploring the biophysics of transcription factor DNA specificity, and the ability to engineer ZF TFs makes them useful tools in biotechnology and synthetic biology. But even with the most recent datasets and models^16-24^, engineering novel ZF TFs with precise sequence specificities remains a challenging problem. The Zinc Finger Consortium’s online database of ZFs^25^ is in principle a useful resource of recognition helices (RHs) that supposedly bind a particular DNA triplet, but characterization assays vary and are relatively incomparable in terms of measuring DNA target specificity and affinity, leading to incongruities in ZF studies^26,27^.

**Figure 1:**
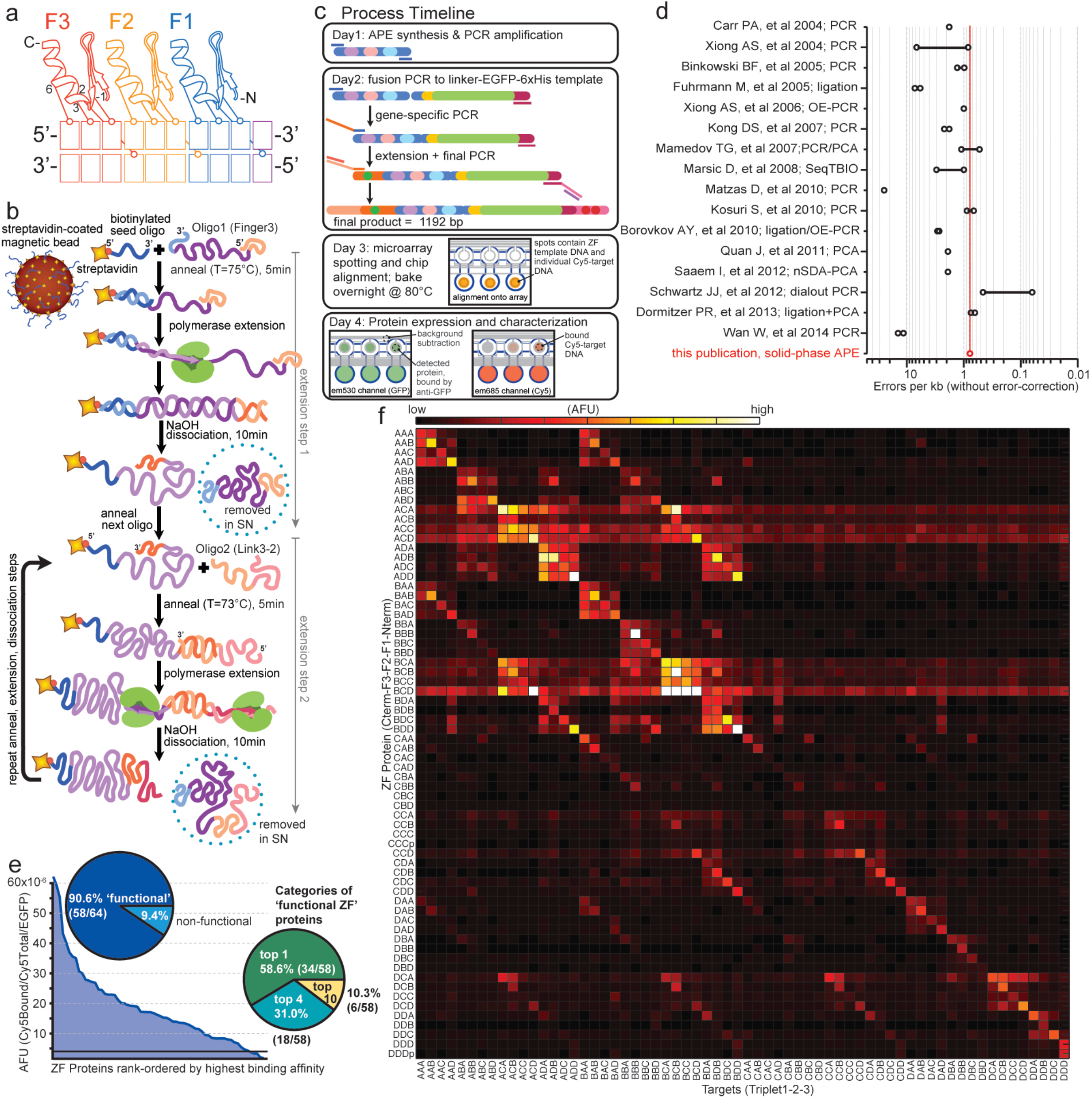
APE-MITOMI applied to ZF TF module combinatorics. (a) Cartoon model of canonical Cys_2_His_2_ ZF TF binding to DNA with residues -1, 2, 3 and 6 of the recognition helix primarily encoding DNA specificity. Residue 2 makes a cross-strand contact, which creates ‘context dependent’ effects. (b) Schematic of the APE solid-phase gene assembly technique, showing assembly through the first two extension steps. (c) Process timeline from gene assembly to protein characterization. (d) Comparison of APE error rate with values from previously published gene assembly techniques. A line between two points indicates a range of error rates from different experimental conditions. (e) Overview of experimental results obtained from combinatoric assembly of ZF TFs demonstrating protein expression and functional DNA binding success rates. (f) Heatmap of dimensionless normalized binding affinities for each assembled ZF TF (y-axis) to 64 predicted consensus DNA targets (x-axis). Protein naming convention indicates ZF domain from C-to-N (F3 to F1), where AAA (A_f3_A_f2_A_f1_)=Zif268, BBB=37-12, CCC=92-1, DDD=158-2 (ref 14); for example, protein ABC = F3 from zif268, F2 from 37-12, F1 from 92-1; target ABC = Zif268 F3 binding consensus triplet GCG, 37-12 F2 binding consensus GAC, 92-1 F1 binding consensus triplet GCC (5’-GCG GAC GCC). Complete target sequences are given in Supplementary Tables.

Gene synthesis by APE (Fig. 1b) is a restriction enzyme-and ligase-free method that operates without the need for transformation, sequencing, and PCR/oligomer purification steps. By coupling APE with cell-free protein expression and MITOMI we reduce the time between protein design and characterization to 3-4 days per iteration cycle (Fig. 1c). For ZF TF engineering, this pipeline consists of 4 stages: (i) solid-phase gene assembly from oligonucleotides, (ii) fusion and overhang extension PCR to generate expression ready linear templates, (iii) microfluidic device programming, and (iv) cell-free protein expression, purification, and DNA-binding characterization (Fig. 1c).

APE is a solid phase synthesis approach based on asymmetric primer extension (Fig. 1b). We succeeded in performing up to 9 consecutive rounds of extension (Supplementary Fig. 1), whereas previous methods achieved up to 4 consecutive extension steps^28^. Gene assembly on magnetic beads allows pooling of identical reactions before fractioning to incorporate unique oligomers in subsequent extension cycles, and sequential addition of each oligomer reduces the risk of forming chimeric products that result from one-pot assembly reactions. In this study we implemented APE assembly using manual bench-top methods but the method is compatible with high-throughput liquid-handling robotic platforms. Our APE gene synthesis exhibited a low error rate of 0.78 errors/kb, which is as good or better than other current gene synthesis approaches based on ligation or PCR assembly (Fig. 1d; Supplementary Tables). For designed ZF TFs, five DNA oligonucleotides are required to assemble the ZF ‘core’ (239 bp), with lengths between 60 and 77nt, and overlaps of 25-28 bp (see Online Methods, and Supplementary Fig. 2) consisting of 3 unique oligomers coding for the three recognition helices, and 2 universal “linking” oligomers.

We first synthesized 64 different ZF TFs and quantified binding of each against a library of the corresponding 64 predicted consensus DNA targets (Fig. 1e,f). APE-MITOMI successfully expressed all 64 ZFs and 90.6% (58/64) of these bound DNA (Fig. 1e). Of those ZFs that bound DNA, 89.7% (52/58) bound the expected consensus target within the top 4 highest-affinity targets and 58.6% (34/58) bound the expected target with highest affinity. To verify that failure to bind and that sequence specificity was unaffected by our approach we cloned and sequence verified the CCC and DDD ZF TF variants generating the plasmid based versions CCCp and DDDp. Both plasmid-based ZF TFs gave identical results when compared to APE generated variants. The heat map in Figure 1f shows the quantitative binding specificity of each of the 64+2 ZF variants. This approach can be used to rapidly identify orthogonal ZF TFs for use in the design and implementation of synthetic genetic networks^14^ and to eliminate ZFs with sub-optimal binding characteristics such as low affinity or extensive non-specific binding. For example, we observed that ZFs containing a ‘C’ finger in position F2 exhibited high levels of non-specific binding, especially when combined with finger variants A and B in position F3.

We next applied APE-MITOMI to engineering ZF specificity. We decided to engineer a ZF with a consensus sequence of ‘GTA GAT GGC’, taking advantage of the relatively well-populated selection of GNN-binding RHs in the Zinc Finger Consortium database (Fig. 2a). We first modified F2 of Zif268 by replacing it with 16 RHs (Fig. 2a and Supplementary Fig. 3) listed as ‘GAT’ binders. Characterizing these variants showed that there was considerable variability in the specificity and binding affinity of the 16 RHs tested. One RH, LLHNLTR, exhibited the highest affinity and specificity for ‘GAT’, and we thus selected this helix for the next step, which involved substituting RH variants into the F1 position to target ‘GGC’. Selecting a helix with good specificity from these RHs proved challenging since nearly all RHs preferred ‘GTC’ and exhibited considerable non-specific binding (Supplementary Fig. 4). A similar specificity profile was observed when we tested the highest affinity F1 RHs in a different background, which reduced the non-specific background (Supplementary Fig. 5). We selected ESSKLKR as the best ‘GGC’ binder in position F1 considering its level of specificity for ‘GGC’ relative to ‘GTC’. We then substituted F3 with RHs reported to bind ‘GTA’. The RH that displayed the highest ‘GTA’ binding affinity and specificity, QSSALTR, was generated by selecting the highest frequency residue in each position derived from all ‘GTA’ RH variants listed in the database (Supplementary Fig. 6). In order to characterize the actual specificity of our synthetic ZF TF (F1 ESSKLKR, F2 LLHNLTR, F3 QSSALTR), a 1-off target library was prepared for the consensus sequences: ‘GTA GAT GGC’ and ‘GTA GAT GTC’. Given the relatively poor performance observed during the F1 selection step, the engineered ZF TF exhibited a surprisingly high affinity and specificity for the intended target consensus sequence (‘GTAGATGGC’). In an attempt to improve the specificity for the intended target, we performed an additional screen for designed RH variants which were anticipated to bind ‘GGC’ using online DNA binding site predictors^20-23^ (Supplementary Fig. 7), but nearly all these variants displayed a higher affinity for GTC and thus failed to further improve the specificity of the engineered ZF TF (Supplementary Fig. 8).

**Figure 2:**
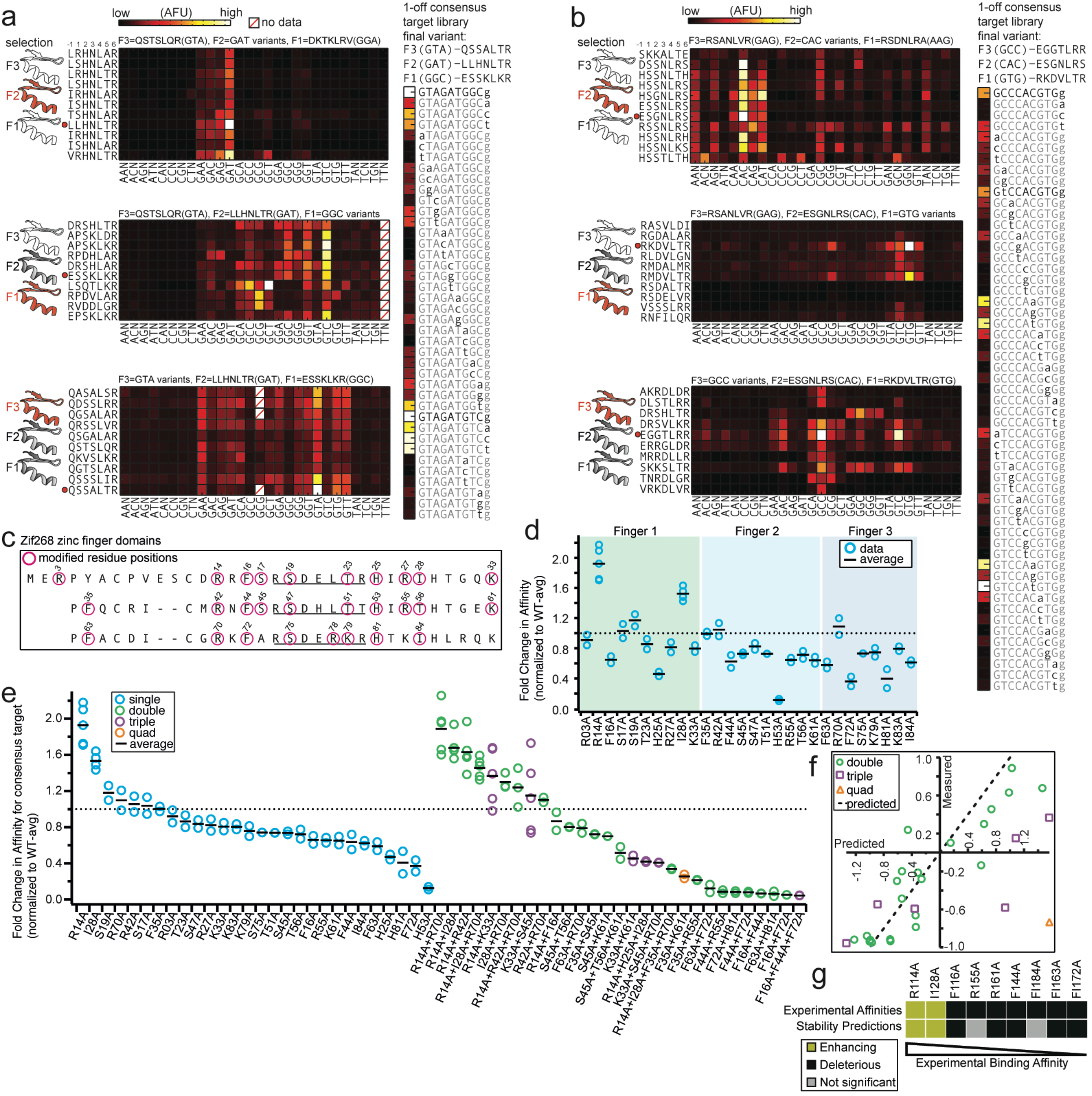
Engineering ZF TF specificity and affinity. (a, b) To the left of each heatmap is a cartoon of the 3 ZF domains in the TF, where ‘hollow’ ZF domains indicate the location of unaltered RHs, red ZF domains indicate the location of the RH variants being varied, and gray ZF domains indicate locations where RHs have already been screened. RH amino acid variants are listed and a red dot indicates the RH that was chosen and used in the subsequent selection round. The vertically-oriented heat map gives the results of the 1-off DNA consensus target library with the final ZF TF design. (a) RH variant characterization across three rounds of screening to engineer a ZF TF that binds the sequence 5’-GTA GAT GGC. (b) RH variant characterization across three rounds of screening to engineer a ZF TF that binds the sequence 5’-GCC CAC GTG. (c) Amino acid alignment of the ZF domains in Zif268 where magenta circles indicate residues that were changed to alanine. (d) Results of single residue alanine scan at the residue positions listed and the resulting fold-change in binding affinity relative the WT zif268 consensus target. (e) Rank ordered fold-change in binding affinity relative to WT consensus target for single and multiple mutant variants. (f) Comparison of measured versus predicted fold-change in binding affinity to determine if a simple additive model can explain the affinity of multiple mutations. (g) Comparison between experimentally measured Zif268-DNA affinities and *in silico* predicted protein stabilities. The subset of mutants with the most pronounced binding changes (excluding histidine to alanine mutants) are shown. Rosetta energies were only considered significant if the absolute changes were above 0.75.

We repeated the same sequential selection process to engineer a ZF TF recognizing the sequence ‘GCC CAC GTG’, which is a more challenging target sequence given the ‘CAC’ in the F2 position (Fig. 2b and Supplementary Fig. 9-11). A number of RHs in positions F2 and F1 were found that bound ‘CAC’ and ‘GTG’, respectively. Finding a high affinity, yet specific RH in position F3 for GCC was more challenging and we had to settle for a RH that bound both ‘GCC’ and ‘GTC’ (Supplementary Fig. 12). Although each individual RH appeared to bind the intended target sequence fairly specifically, when testing the assembled ZF TF using a one-off library the consensus target was ‘GTC CAT GTG’, but ‘GCC CAC GTG’ could also be bound albeit with a lower affinity. Other RHs from the initial ‘GCC’ screen, which displayed lower affinity but higher specificity for ‘GCC’ were also tested against a one-off library (Supplementary Fig. 13), but all of these exhibited higher affinity for other targets than ‘GCC CAC GTG’. These results indicate that engineering ZF TFs to bind a particular sequence is relatively easy to achieve using a stepwise selection process, but engineering the precise specificity landscape of a ZF TF is more challenging. The individual specificity profiles of RHs can vary considerably, and in some instances do not reflect their annotated sequence specificity given in the ZF Consortium Database or online binding site predictors. One significant advantage of APE-MITOMI based assembly and characterization is the fact that the method returns information on the precise specificity landscape and affinity of the synthesized transcription factors during all stages of assembly. APE-MITOMI allows the generation of TFs with similar consensus sequences but different specificity and/or affinity profiles and enables the examination of these characteristics for the function of native and synthetic transcriptional regulatory networks.

Focus in ZF engineering has been on rationally designing ZF specificity, but it is unclear to what extent affinity can be tuned and whether affinity can be tuned independently of sequence specificity. The ability to tune ZF affinity is of interest in creating synthetic transcriptional regulatory networks^14^ and it would drastically simplify the task of engineering ZF TFs if affinity could be tuned independently of specificity. Based on the Zif268 crystal structure we selected 28 residues that could be involved in determining the protein’s affinity to DNA, and changed these residues to alanine (Fig. 2c). As expected, most modifications resulted in modest decreases in affinity as compared to wildtype (WT) Zif268 (up to 2-fold). To our surprise, a few modifications led to increased affinity (R14A, I28A, R70A) (Fig. 2d). Given that the single substitutions resulted in modest affinity decreases, we tested 22 double substitutions, 6 triple substitutions and 1 quadruple substitution (Fig. 2e). These novel mutants allowed us to extend the dynamic range to 1/5 of WT and allowed us to smoothly tune affinity over the entire accessible range between 2x increased and 5x reduced affinity. We could show that most of the affinity mutants retain their specificity landscape, indicating that affinity can be tuned independently of specificity (Supplementary Fig. 14). We also tested whether a simple additive model could predict double, triple and quadruple substitutions (Fig. 2f). Overall, the simple additive model captures the trend for double substitutions but performs poorly for triple and quadruple substitutions.

Next, we sought to understand whether the observed differences in affinity could be predicted by energy calculations performed *in silico* based on the available structural information for Zif268. To do so, we modeled single alanine mutants with the Rosetta protein modeling software^29^. First, we analyzed the effect of the alanine mutants on the Zif268-DNA binding energy and perhaps not surprisingly, given the low number of atomic interactions established across the interface for the large majority of these residues, we did not find any large differences in the binding energies of the point mutants in comparison with the WT Zif268. Bolstered by the differences in affinity observed in mutants of zinc-coordinating residues (H25A, H53A and H81A), which recognizably play a critical role for structural integrity of the zinc finger fold^30^, we asked if other mutations could also cause considerable losses in protein stability and consequent losses in DNA binding affinity. To address this question we performed *in silico* δδG calculations using Rosetta. Being aware of the limitations of current modeling techniques in accurately describing subtle structural and functional effects, we focused on mutants that exhibited pronounced differences in binding (reductions < 0.7 and improvements > 1.3 – 9 in total). In this subset of mutants (7 deleterious and 2 enhancing; Fig. 2g) we were able to correctly predict the variations in binding affinity with our stability predictions for 6 of the 9 mutant proteins, strongly suggesting that the observed binding affinity differences are linked to protein stability.

We coupled a gene synthesis strategy directly with high-throughput on-chip protein expression, purification, and quantitative characterization, showing that it is possible to completely circumvent molecular cloning and cell-based processing steps generally needed in protein biochemical analysis. Integration of these methods reduced the time required between protein design and protein characterization to 3-4 days. In this study we synthesized over 440 ZF TF variants, and characterized the binding specificity of each for a total of over 98,000 measurements. By comparison, the human genome encodes ∼675 C_2_H_2_ ZF TFs^31^. We have previously shown that over 400 full-length drosophila transcription factors could be expressed on a single MITOMI device^32^ indicating that there is no major bottleneck to on-chip protein expression. Although we synthesized all 440+ genes in this study by hand, the APE gene synthesis method can be readily automated on standard liquid handling robots eliminating this last bottleneck in the process. It should thus be possible to characterize thousands of synthetic protein variants per week, which in turn will enable exploration of the protein sequence-function relationship in unprecedented detail, aid in the development of accurate computational predictions of protein function, and allow us to rapidly engineer novel proteins.

## Acknowledgments

We thank members of the Maerkl Lab for helpful discussions, the Deplancke Lab for use of their ArrayWoRx scanner, and EPFL for financial support.

## Author contributions

MCB and SJM conceived of the synthesis technique, designed the project, and analyzed the data. MCB and EP tested early protein variants, optimized the gene synthesis method and determined the error rate. MCB prepared all of the ZF TF templates and conducted all of the MITOMI experiments. BEC performed the *in silico* analysis. MCB, EP, BEC, and SJM wrote the paper.

## ONLINE METHODS

### I. Gene Synthesis Protocol

#### Synthetic DNA Design

All zinc finger modules used in this study were either taken from the Zinc Finger Consortium’s online Zinc Finger Database^25^, or from references 14 and 22, which contain natural and designed single zinc finger modules or concatenated, multi-finger arrays. Other zinc finger modules were designed using online DNA binding predictors^20-23^. For our purposes, we use only the 7 amino acid residues in positions -1 to 6 of the α-helix. Other parts of the protein that are indirectly involved in DNA sequence recognition, called the framework, are taken directly from the three-finger murine transcription factor Zif268 (RCSB PBD 1AAY; amino acid sequence: MERPYACPVESCDRRFS<u>RSDELTR</u>HIRIHTGQKPFQCRICMRNFS<u>RSDHLTT</u>HIRTHTGEKP FACDICGRKFA<u>RSDERKR</u>HTKIHLRQKD, where underlined residues are the locations of the native first, second and third α-helices). The Zif268 coding sequence (90 aa) was converted to an *E. coli* (strain K12) codon optimized nucleotide sequence using JCat (Java Codon Adaptation Tool^33^, www.jcat.de). The resulting 270 nt sequence was then partitioned into 5 oligomers: 3 ‘finger’ oligomers containing the sequence coding for α-helix residues with flanking sequence, and 2 ‘linking’ oligomers containing sequences bridging the three ‘finger’ oligomers (see Supplementary Fig. 2 and Supplementary Tables for sequences). Each oligomer has a 25 or 28 nt overlap with the oligomer preceding it for annealing. This way, all oligomers with the appropriate flanking sequence can be interchanged with each other since they contain the necessary complementary sequence for annealing. This allows single oligomers to be used in multiple zinc finger assemblies, but limits them to the same position in the assembly process. Linking oligomers ‘Link3-2’ and ‘Link2-1’ are used for all assemblies, whereas libraries of O1F3 (Oligo1 Finger3), O3F2 (Oligo3 Finger2) and O5F1 (Oligo5 Finger1) oligos are used to generate different 3-finger assemblies (21 nt located in the colored regions of Supplementary Fig. 2). Oligomers were ordered from IDT with standard desalting only and were rehydrated to 500 μM in 1x Tris-EDTA (TE) buffer for stock solutions.

#### Asymmetric primer extension (APE) assembly

Prior to gene-assembly, an aliquot (800-1200 μL) of MyOne™ Streptavidin T1 beads (Life Technologies) are placed in a 1.5 mL Eppendorf tube, pelleted using a magnetic stand, and resuspended in an equivalent volume of 0.2M NaOH in water. The beads are preconditioned for at least 1 hour at room temperature before use, then stored at 4°C for longer conditioning times. These conditioned beads can be used for up to one month after being suspended in the 0.2M NaOH solution. It was found that preconditioning of beads in NaOH releases labile streptavidin monomers^34^ and in our hands this translated to a reduction in non-specific PCR bands during intermediate quality control PCR steps and during amplification of the final assembly. Each individual assembly reaction requires 25 μL of preconditioned beads. Lower bead quantities may work as well, but to account for losses during washing and transfer steps, we have continued using 25 μL with consistent success. Larger reactions are also possible by scaling all volumes accordingly. This is particularly useful during the creation of zinc finger combinatoric array variants. By starting with a large pool of beads, beginning the assembly together with the same Oligo1Fin3 and Link3-2, followed by partitioning the pool into smaller volumes and continuing assembly with different Oligo3Fin2 parts in separated reactions, followed by a final partitioning for Oligo5Fin1 parts, where the final volume in each Oligo5Fin1 assembly is 25 μL, many different genes can be assembled within the same workflow.

For a single APE reaction, 25 μL of preconditioned beads are pelleted using a magnetic stand (Invitrogen DynaMag™-Spin) for 30-60s until the solution is clear. The supernatant (0.2M NaOH) is carefully aspirated. The beads are then washed twice with 25 μL of 1x binding and washing buffer containing Tween20 (B&W+Tween) (each washing step involves adding the wash solution, mixing the solution by aspiration until the beads are resuspended, then pelleting the beads and removing the supernatant), then pelleted again and resuspended in 25 μL of 2x BW Buffer, to which 25 μL of ‘seed’ oligomer solution (0.12 μM biotinylated initiation oligomer in PCR grade water) is added and mixed. This mixture is incubated at room temperature for at least 15 minutes on a lab rotisserie. Following incubation, the beads are pelleted against the magnetic stand, washed twice with 25 μL of 1× HF Buffer without detergent (New England Biolabs) to prevent bubble formation during resuspension, and finally resuspended in 25 μL Oligo1Fin3 extension mix (final concentrations: 1x HF Buffer with detergent, 0.2 mM dNTPs, 5% DMSO, 8 μM Oligo1Fin3, 0.3 units Phusion High-Fidelity Polymerase (NEB)). This mixture is then placed on a thermocycler and run through a brief annealing and extension routine (5.5min at T_anneal_, 2min at 72°C, then hold at 25°C; T_anneal_ for each oligomer is given in Supplementary Fig. 2). The tube is removed from the thermocycler, the beads are pelleted, and the supernatant is removed and discarded. The beads are then washed twice with 50 μL 1× SSC buffer, resuspended in 50 μL 0.15M NaOH and incubated at room temperature on a rotisserie for 10 minutes to facilitate strand dissociation. The beads are then pelleted, the supernatant is removed, and the beads are washed once with 50 μL 0.15M NaOH, once with 50 μL 1× BW+Tween, and once with 50 μL 1× HF buffer without detergent. The beads are then resuspended in Oligo2 (Link3-2) extension mix (same recipe as for Oligo1, except Link3-2 is used), then placed back on the thermocycler, and run through the annealing and extension routine, where T_anneal_ has been adjusted to the temperature required for this annealing reaction. This procedure of extension, strand dissociation via 0.15 M NaOH, and buffer exchanges is repeated for each oligomer in the assembly. After the final extension reaction (Oligo5Fin1), there is no NaOH dissociation step. Instead, the beads are pelleted, washed twice with 50 μL 1× SSC buffer, and resuspended in a final volume of 20 μL 10 mM Tris-Cl pH 8.5 buffer.

The beads in Tris-Cl buffer from the final extension step are used directly as template for a PCR amplification of the complete 5-step assembly product. PCR primers were designed to amplify the 239 bp product. For a 20 μL PCR reaction, final concentrations are as follows: 1x HF Buffer with detergent (New England Biolabs), 0.2mM dNTPs, 5% DMSO, 0.5 μM each primer (assembly_check-f and –r), 0.6μL suspension of beads in Tris-Cl (template), 0.3 units Phusion High-Fidelity Polymerase; touchdown PCR: 98°C, 30s; 74>72°C, 30s then 17 cycles at 71°C, 30s ; 72°C, 30s). 4 μL of this PCR is then run on a 2% agarose gel with 0.4x GelGreen (Biotium) at 110V for 1h. Due to non-specific primer interactions with the template and interactions with truncated assembly products, some PCR amplifications can result in the formation of multiple bands, the largest of which is the complete assembly product. To overcome this problem, we have taken advantage of a modified band-stab technique^35^ to isolate the band of interest and re-amplifying it to reduce the amount of nonspecific products in downstream steps. Briefly, the 2% gel is imaged using a blue-light transilluminator and the band of interest is captured using a 200 μL pipette tip, with the end cut off about 1 cm from the tip, by stabbing into the gel at the location of the band. The pipette tip is then placed into a 1 mL Eppendorf tube, and the agarose gel core inside the pipette tip is pushed out using a second sterile pipette tip. 20 μL of Qiagen EB buffer (10 mM Tris-Cl, pH 8.5) is added to the agarose gel sample, briefly vortexed, centrifuged, and incubated at 80°C for at least 10 minutes with the tube cap closed. The sample is then vortexed and centrifuged again, before a 0.25 μL sample of the buffer is taken as template for a second PCR amplification. This PCR is prepared and thermocycled following the same recipe from the first PCR (assembly check PCR; Supplementary Fig. 15).

Following the second assembly check PCR, the core region coding for the three linked a-helices is complete, but the final template will consist of a C-terminal proline-linker and EGFP fusion, a 6x histidine tag, and 5’ and 3’ UTRs for expression within a cell-free, transcription/translation mixture. All of these parts are added to the zinc-finger assembly via 4 different PCRs: a fusion PCR (for adding the proline-linker, EGFP domain, and 6×-histidine tag; results in a 1019bp product), a gene-specific PCR (for adding part of the 5’ and 3’ UTRs; results in a 1084bp product), and an extension+final 2-step PCR (which completes the template construction and amplifies the full-length product of 1192bp; Supplementary Tables).

The fusion PCR requires two templates: the 239bp zinc-finger construct and a previously amplified EGFP domain from the pKT127 plasmid, including a 5’-proline linker and 3’-6x histidine tag. Briefly, in a 20 μL PCR containing 1× HF Buffer with detergent, 0.2 mM dNTPs, 5% DMSO, 0.5 μM each primer (Prolinker-EGFP-f and EGFP-6His-r), 1 ng of pKT127, and 0.3 units Phusion High-Fidelity Polymerase are thermocycled for 25 cycles (98°C, 30s; 61.7°C, 30s; 72°C, 1min). This product (Prolink-EGFP-6His, 805nt) is used without purification in the fusion PCR. The fusion PCR is carried out in 2 steps, the first reaction contains all the necessary ingredients for PCR except the nucleotide mix and polymerase. A 15 μL reaction is prepared containing 1x HF Buffer with detergent, 5% DMSO, 0.5 μM of each primer (assembly_check-f and EGFP-6His-r), and 0.25 μL each of the assembly PCR from band-stab and Prolink-EGFP-6His PCR. This mixture is placed on a thermocycler and heated to 98°C for 4min, then cooled down (10% ramp) to 25°C for annealing. Then 5 μL of an extension mixture (1x HF Buffer with detergent, 0.2 mM dNTPs, 0.3 units Phusion) is spiked into the annealing mixture (20 μL total) and cycled 20 times (98°C, 30s; 72°C, 30s; 72°C, 1min).

The gene-specific PCR uses the product generated in the fusion PCR as template. A 20 μL reaction is prepared: 1x HF Buffer with detergent, 0.2 mM dNTPs, 5% DMSO, 0.5 μM each primer (genespec-f and EGFP-6His-r), 0.25 μL of the fusion PCR, and 0.3 units Phusion polymerase. This reaction is cycled using a short touchdown PCR (98°C, 30s; 75>72°C, 30s; 72°C, 1min), followed by 16 cycles (98°C, 30s; 72°C, 30s; 72°C, 1min).

The 2-step extension+final PCR uses the product generated in the gene-specific PCR as template. In this reaction, it is very important to use the HF Buffer without detergent, since this product will be used directly for microarray spotting and the presence of detergent will result in large spots. A 20 μL reaction is prepared: 1× HF Buffer without detergent, 0.2 mM dNTPs, 5% DMSO, 2.5 nM each primer (ext-f and ext-r), 0.25 μL of 1:10 diluted gene-specific PCR (in Tris-Cl or water), and 0.3 units Phusion polymerase. This mixture is thermocycled 10 times (98°C, 30s; 61°C, 30s; 72°C, 1min). Then the reaction is kept at 72°C for 2 minutes, and cooled to 25°C. At this point, 0.1 μL of each final_highTm primer (50 μM stock; final 0.25 μM in 20.2 μL) are spiked into the mixture, and it is thermocycled again via a short touchdown PCR, 98°C, 30s; 75>72°C, 30s; 72°C, 1min, then 20 cycles 98°C, 30s; 71°C, 30s; 72°C, 1min. Successful amplification is determined by running 1.5 μL of the product on a 1% agarose gel, and checking for the 1192bp product (see Supp Fig 15).

Materials:

a. Dynabead MyOne Streptavidin T1, preconditioned in 20mM NaOH solution*
b. Magnetic stand for attracting magnetic beads
c. PCR tubes with attached flat caps (VWR, 300 μL)
d. Lab rotisserie
e. Thermocycler
f. Microcentrifuge tubes with attached caps (Eppendorf, 1 ml)
g. B&W (Binding and Washing) Buffer for Dynabeads (2x : 10mM Tris-HCl pH 7.5; 1mM EDTA; 2M NaCl)
h. B&W + Tween20 Buffer – take one volume of B&W buffer (2x) and dilute with one volume of 0.01% Tween20 in DI water**
i. NEBuffer 2 (B7002S, 10x) (1x : 50mM NaCl, 10mM Tris-HCl, 10mM MgCl_2_, 1mM DTT, pH 7.9 @ 25°C)
j. Phusion High-Fidelity PCR kit (NEB, E0553L, 200rxn)
k. Phusion® HF Buffer Detergent-free (5x) (NEB)
l. DMSO (100%)
m. Synthesized DNA Oligomers (IDT, varying lengths), rehydrated in 1x TE buffer to 500uM stock solution, then diluted to 50μM for working solution
n. Deoxynucleotide (dNTP) Solution Mix (NEB, N0447S, 10mM each)
o. SSC Buffer 20x Concentrate (Sigma, S6639-1L) – dilute to 1x in PCR-grade water
p. Tween20 for molecular biology (Sigma, P9416-50mL) – prepare 0.01% Tween20 in PCR-grade water
q. 2M NaOH (prepare and store with parafilm-wrapped cap, water vapor from the air can be absorbed)*** – dilute to 0.15 M in PCR-grade water, prepared fresh on day of synthesis reactions

#### Footnotes

* preconditioning of beads in NaOH releases labile streptavidin monomers, precondition for at least 1 hour at room temperature before using, store @ 4°C for longer conditioning times^34^

** use of 0.01% v/v Tween20 in buffers to reduce non-specific binding (Dynabeads manual, and ref 36)

*** wear safety goggles to prevent splashing in eyes, highly caustic/strong base, causes burns

#### Error Rate Analysis

Gene synthesis reaction on beads was carried out using either DNA Polymerase I, Large (Klenow) Fragment (NEB) for APE assembly reactions entirely carried out at room temperature, or Phusion High-Fidelity Polymerase with brief annealing and extension steps on a thermal cycler. Both approaches used unpurified oligomers for the construction of zinc finger array 92-1 within a Zif268 backbone (Supplementary Tables).

##### Klenow assembly

Following 20 cycles of PCR using Phusion polymerase for amplification of template detached from beads using an SDS-boiling and reannealing protocol, the reaction was run on an agarose gel and the product band was gel-stabbed, and a second PCR (20 cycles) was run.

##### Phusion assembly

Following 20 cycles of PCR using Phusion polymerase for amplification of the template attached to beads in 1x SSC buffer, the reaction was run on an agarose gel and the product band was gel-stabbed, and a second PCR (20 cycles) was run.

This PCR product (239bp) from each of the band-stab PCRs was purified and cloned via Gibson assembly into the pUC19 plasmid with assembly-check overhangs. In addition, the PCR from the Phusion assembly was used without the band-stab procedure, to determine whether the band-stab has an effect on error rate.

Chemically competent DH5α *E. coli* cell aliquots (30 μL) were transformed with 1.5 μL of each Gibson assembly product via heat shock (30s at 42°C), recovered in 300μl SOC medium for 1 hour at 37°C, and plated on ampicillin plates for overnight growth at 37°C. Colonies from each plate (Klenow+band-stab, Phusion+band-stab, Phusion no stab) were picked with a sterile 200 μL pipette tip, briefly stirred in 20 μL PCR-grade water, boiled for 15min, and centrifuged. Water from the colony boils was used as template for an insert-check PCR using the primers pUC19-f and pUC19-r with Phusion polymerase. All of these PCR reactions were run on 1.5% agarose gels with GelRed to determine which colonies had the correct sized insert. Colonies were picked and analyzed in this way until 32 colony PCRs for each assembly method were identified with a single band corresponding to the correct sized insert. The insert-check PCRs were submitted for Sanger sequencing in a 96-well plate without PCR cleanup (Microsynth AG), and the resulting sequencing reads were aligned with the expected sequence to analyze the error-rate and identify which types of errors were prevalent.

The highest error rate was observed in the colonies cloned with the Klenow APE assembly method, however the majority of these products would have resulted in nonfunctional ZF TFs since they code for early stop codons or cause frame-shifts which would prevent the EGFP tag from folding and being captured in a MITOMI experiment. The Phusion APE assembly products yielded significantly lower erroneous sequences, with no effect seen from the band-stab PCR. The high error rate seen in the Klenow reaction product can be partly explained by comparing the relative error rates documented for Klenow and Phusion polymerase.

### II. dsDNA Target Synthesis

Double stranded DNA targets (dsDNA) for zinc-finger array binding were prepared via isothermal Klenow extension as previously described^32^. Oligomers were designed such that the DNA contains the 9 nt target sequence with single nucleotide flanks (11 nt total). At the 3’end of the DNA is the complementary sequence for a Cy5-labeled primer (5’CompCy5). The reaction consists of 2 steps, one annealing step, and one extension step. Each 20 μL annealing reaction contains 1x NEB Buffer 2, 10 μM 5’CompCy5, and 15 μM Target oligomer. This mixture was placed on a thermocycler, heated to 94°C for 5 minutes, then cooled to 37°C (10% ramp) for 5 minutes, then held at 20°C. Following the annealing program, 10 μL of extension mix (1x NEB Buffer 2, 3mM dNTPs, and 2.5 units Klenow exo-) are spiked into the annealing mix, and the reaction is thermocycled according to the following routine: 37°C, 90min; 75°C heat kill, 20min; 10% ramp down to 30°C, 30s; hold at 4°C.

### III. Microarraying

#### Preparation of epoxy-silane glass slides

For microarray printing, epoxy-silane functionalized glass slides were prepared, following an adapted protocol^37^. A MilliQ water and ammonia solution (NH_4_OH, 25%) mixture was prepared in a 5:1 ratio, respectively, and heated to 80°C. Then, 150 mL of hydrogen peroxide (H_2_O_2_, 30%) was added to the mixture, and glass microscope slides were placed in the cleaning bath for 30 minutes. The glass slides were then removed and rinsed in fresh MilliQ water, dried with N_2_, and placed in a second bath containing 1% 3-glycidoxypropyltrimethoxymethylsilane (97% purity) in toluene, and incubated for at least 20 minutes at ambient temperature. Then, the glass slides were removed, rinsed in fresh toluene, dried with N_2_, and baked at 80°C for 30 minutes. The glass slides were removed from the oven, allowed to cool, and stored under vacuum at room temperature in opaque storage boxes until used. Immediately prior to microarray printing, both sides of each glass slide is briefly rinsed with fresh isopropanol, dried with N_2_, rinsed with fresh toluene, and dried with N_2_ again.

#### Sample microarraying

All samples to be printed onto a microarray were prepared in a 384-well microtiter plate. Each zinc-finger array assembly (1192 bp PCR product) was co-spotted with a target DNA (Klenow extended products) in duplicate onto epoxy-silane coated glass slides using a microarray robot (QArray2) with a 946MP4 microspotting pin (Arrayit). Up to four slides were prepared in a single printing session. Each spot on the array was generated by four consecutive printing programs. Immediately following completion of printing from a given sample plate, the sample wells were covered with adhesive PCR foil seals (ThermoScientific) and stored at -20°C until needed for future printing runs. In general, Klenow target plates were re-used for up to 5 printing runs before the volume of each well became too low to be used. After the final printing program, the microspotting pin is cleaned by sonication for 15min in a 15 mL Falcon tube containing one drop of dish detergent and 10 mL of deionized water, then sonicated for 30min in a 15 mL Falcon tube containing 10 mL of 70% ethanol in water. Before use, the pin tip is rinsed under deionized water, dried using a high pressure (100 psi) compressed air gun, then inspected under a microscope to verify the tip is clean and free of debris. Finally, the shaft of the pin is briefly polished with a dry Kimwipe to prevent sticking when placed into the robotic printer head.

#### Printing Routine

1. 0.5% BSA in water prespot, 1 stamp/spot, 4 stamps/ink, 1 well per template; 45% humidity, no same sample wash
2. template spotting (6 μL PCR product + 64 μL 2% BSA in water), 2 stamps / spot, 4 stamps/ink; 45% humidity, wash every 32 inks
3. 0.5% BSA in water overspot, 1 stamp / spot, 4 stamps / ink, 1 well per template; 45% humidity, no same sample wash (use same wells from ‘prespot’)
4. dsDNA target spotting (10 μL Klenow product + 60 μL 2% BSA in water), 3 stamps / spot, 3 stamps / ink; 45% humidity, wash after 32 inks

The first BSA printing routine is to block the surface of the glass, to limit the amount of template and target DNA that is irreversibly attached to the epoxy-silane surface. The second BSA printing routine is implemented to deposit a thin blocking layer on top of the printed template spots, to reduce the risk of cross-contamination due to template carry-over during target printing (since target printing does not involve same-sample washing steps).

### IV. MITOMI chip fabrication and operation

#### Mold Fabrication

The MITOMI microfluidic device^6,7^ (Supplementary Fig. 16) consists of two superimposed layers, the flow layer and the control layer. Each layer is fabricated in polydimethylsiloxane (PDMS) using 4” silicon wafer molds fabricated using standard lithography techniques^38^. Each wafer (control and flow) contained three pattern replicates for a 1024-chamber (16 rows by 64 columns) MITOMI device.

Mask fabrication was carried out using a Heidelberg DWL200 laser lithography system with 10mm writing head and solid state wavelength stabilized laser diode (max. 110 mW at 405 nm). Each layer of the MITOMI device was reproduced as a chrome mask. After laser writing, the chrome mask is cycled twice for 15s in developer mixture (1:5 MP351 and deionized water, respectively), 45s of agitation, then rinsed and dried. The developed mask is then chrome etched for 110s, rinsed, cleaned twice for 15min in 1165-remover bath, rinsed and air dried.

The flow layer mold is first cleaned for 7 min in a Tepla300 plasma stripper with 400 mL/min O2 at 500W and 2.45 GHz. The wafer is then treated with hexamethyldisilazane (HMDS) using an ATMsse hotplate at 125°C for 12min. Positive photoresist AZ9260 is spin-coated on the cleaned wafer for 10s at 800 rpm, then 40s at 1800 rpm (ramp 1000 rpm/s) to produce a substrate height of 14 μm. The wafer is then baked on a 115°C hotplate for 6min. The soft-baked positive resist is then allowed to rehydrate for 1h. The wafer is then exposed during 3 intervals of 18s with a 10s pauses between each exposure on a MA6 mask aligner (power 360 mJ/cm^2^, intensity 10 mW/cm^2^, broad-spectrum lamp, hard contact exposure mode). After a 1h relaxation time, the wafer is developed in a DV10 chamber via multiple, automated cycles of rinsing/agitation with development mixture (1:4 ratio of AZ400K and deionized water, respectively) until the features are visible. Finally, the wafer is heated to 160°C for 20 minutes to anneal and round the features of the flow wafer to create a profile that allows complete valve closure.

The control layer mold is first cleaned following the same plasma treatment protocol as the flow layer mold. Negative photoresist SU-8 GM1060 (Gestertec) is spin-coated on the cleaned wafer for 10s at 500 rpm (ramp 100 rpm/s), 10s at 1500 rpm (ramp 100 rpm/s), 1s at 2500 rpm, and finally 6s at 1500 rpm to produce a substrate height of 14 μm. The wafer is baked on a hotplate for 30min at 130°C, then 25min at 30°C. The wafer is then exposed on a MA6 mask aligner for 13.2s (power 360 mJ/cm^2^, intensity 10mW/cm^2^, broad-spectrum lamp, hard contact exposure mode). The exposed wafer is developed manually by bathing in PGMEA twice for 1.5min, then rinsed in isopropanol and dried with an air gun.

#### Device Fabrication

Prior to PDMS casting, both the flow and control layer wafers are subjected to vapor deposition of trimethylchlorosilane (TMCS, EMD Millipore Corp.) for at least 30min by placing them within a sealed plastic container with a small dish containing 0.25 mL liquid TMCS. TMCS treatment is repeated for at least 15min before all subsequent PDMS casting rounds.

The control layer wafer is placed into an aluminum foil-lined glass Petri dish, and 60g of Sylgard elastomer (5:1 mix of elastomer base and curing agent, respectively) is mixed for 1min at 2000rpm (400×g) and degassed for 2min at 2200 rpm (440×g) in a centrifugal mixer. The elastomer mixture is poured on top of the control layer in the Petri dish, and degassed in a vacuum dessicator for 20min at ambient temperature.

For the flow layer, 21 g of PDMS mixture is prepared at the ratio of 20:1 (base:curing agent), then mixed and degassed in a centrifugal mixer according to the same speeds and times as the control layer. The flow wafer is carefully centered on top of a spin-coater platform using wafer tweezers, and the flow layer PDMS mixture is poured in the center, taking care not to create any bubbles. The mixture is spin-coated onto the wafer with a 15s ramp and 35s spin at 2800 rpm.

The degassed PDMS on the control layer wafer is removed from the vacuum chamber. Residual bubbles are removed with a scalpel and any pieces of dust are carefully removed from the control channel grid using the tip of the scalpel blade. Both the control and flow layers are then placed into an oven at 80°C for 28-30min. After baking, both Petri dishes are removed from the oven and briefly allowed to cool. The control layer is then cut with a scalpel in a rectangle around each pattern replicate, and each rectangle of cured PDMS is carefully peeled away from the silicon wafer. Holes are punched through each of the control line input channels on the patterned side of the PDMS block. The patterned side of the control layer is cleaned twice with Scotch Magic Tape to remove dust and debris then quickly placed on top of the flow layer replicates. A stereomicroscope is used to precisely align the features of the control layer so that they overlap with the chambers visible on the flow layer. Once aligned, the assembled device is bonded at 80°C for 90-180min. The bonded devices are removed from the oven and briefly allowed to cool. A scalpel is guided around the outer edge of the control layer PDMS block to cut the thin flow layer. Then each individual device is gently peeled from the flow layer wafer, and holes are punched through the patterned side inlets and outlet of the flow layer. Each device is then cleaned with Magic Scotch tape and trimmed to fit within the boundary defined by the glass slide-holding cartridge of the microarray scanner. The assembled device is aligned with a printed microarray on an epoxy-silane glass slide using a stereomicroscope and bonded overnight at 80°C before use.

The flow layer mold is cleaned of residual polymerized PDMS by pouring on another layer of mixed PDMS (this can be leftover control-or flow-layer PDMS mixtures prepared earlier; to stall cross-linking for several hours, store the PDMS mixture at 4°C), and baked at 80°C for at least 1 hour. The resulting thicker layer of PDMS can be easily peeled away from the flow layer, resulting in a clean surface to repeat the process. Both the cleaned flow wafer and control wafer are cleaned with high pressure (100 psi) compressed air gun to dislodge pieces of dust or PDMS before being treated with TMCS.

#### Device Setup

Assembled MITOMI chips (Supplementary Fig. 16) bonded to microarray printed glass slides were stored at 40°C following an overnight bonding at 80°C until used. To begin an experiment, control line tubing is filled with PBS using a syringe, and pins are placed into the appropriate locations to feed into the control valves of the microfluidic device. The control lines are actuated at low pressure (10 psi) to begin filling the control lines of the microfluidic device. Once all of the control lines are filled, the sandwich valves and button valves are deactivated, and the pressure is increased to 20-22 psi to ensure complete closure of all other valves. At this point, a 40 cm long line of Tygon® tubing with a blunt-end needle at one end is filled with 50 μL biotin-BSA in deionized water (2 mg/mL), and inserted into the punched hole in the second position of the flow layer inlet tree. The tubing is then connected to a low pressure line (3.5 psi) and the valve controlling the flow of this line is deactivated. Then, the valve controlling the wash-out exit of the inlet tree is deactivated, and biotin-BSA from the inlet well should begin flowing immediately towards the exit well. The exit well valve is then activated, forcing the biotin-BSA to fill the air space in the inlet tree, which prevents the air flowing on towards the main body (16×64 chamber array) of the device. Once all the air is eliminated from the inlet tree, the valve controlling flow onto the chamber array is deactivated, and the biotin-BSA is allowed to flow through the flow space, towards the exit at the end of the device. The exit valve is activated to force the biotin-BSA to fill all of the void space in the flow layer, effectively binding to the surface via the epoxy-silane groups. The exit valve is then deactivated, and biotin-BSA is allowed to wash across the flow layer for 10-15min. After the initial blocking step, another 40cm long line of Tygon® tubing with a blunt-end needle at one end is filled with 75 μL 0.01% Tween20 in PBS. The flow of biotin-BSA is stopped, the Tygon® tubing is disconnected from the low pressure line, and the valve to the chip and the

#### Surface Derivatization, Protein Synthesis, Binding Assay, and Device Readout

Biotin-BSA (2 mg/mL) is flowed through the device for 15min at 3.5 psi. The chip is then washed with 0.01% Tween20 in PBS for 5min to wash away unbound biotin-BSA. Next, neutravidin (1 mg/mL) is flowed for 15min followed by 0.01% Tween20 in PBS for 5min. The button valves are then activated and biotin-BSA is again flowed across the chip for 10min, blocking all of the neutravidin binding sites except those protected under the area of the button valve. The chip is again washed with 0.01% Tween20 in PBS for 5min. Then a solution containing 0.5 μL biotinylated antibody to GFP (1 mg/mL stock, Abcam ab6658) in 100 μL 1% BSA in PBS is flowed across the chip for 5min, the button valve is deactivated, and the antibody solution is flown for 15min, allowing the antibody to bind to the available neutravidin under the button valve. Then, the chip is flushed with 0.01% Tween20 in PBS for 5min, and with PBS for 5min. The button valve is then activated, and ITT mixture (NEB PURExpress, 10 μL SolnA, 7.5 μL SolnB, 0.5 μL RNAse Inhibitor (Roche), 7 μL PCR grade water) is flowed for 10min. The exit valve is activated for 2min while the ITT is being flowed on-chip to build up pressure. The neck valve is deactivated, and the ITT mixture is allowed to fill the DNA chambers for 1-2min. Once the DNA chambers are filled, the neck valves are activated, the exit is opened, and fresh ITT is allowed to flow across the chip for 10min. Then the sandwich valve is activated while flowing ITT mix during the last minute of ITT washing. Once the sandwich valves are partially closed, the button valve is deactivated, the neck valve is deactivated, the exit is closed, and the flow of ITT is stopped. The inlet tree valve controlling entry to the chamber array is closed, and the inlet tree is briefly flushed with 0.01% Tween20 in PBS. Then the entire chip is placed on a flatbed thermal cycler set to 37°C, and incubated for 3-5h. During this time, the DNA array spots are rehydrated in the ITT mix, transcription and translation occur, synthesized zinc-finger/EGFP fusion protein diffuses and is bound by the anti-GFP moiety located under the button valve, and target DNA diffuses and interacts with the various zinc finger DNA binding domains. After incubation, the chip is placed into an ArrayWoRx microarray scanner, and an image is taken in three fluorescent channels (A488/GFP, Cy3, Cy5) to determine relative amounts of solution phase target DNA in the MITOMI chamber. The button valves are then activated, the sandwich valves are deactivated, and the neck valve is activated again. The flow space is washed with 0.01% Tween20 in PBS for 5min to remove unbound target DNA, then the chip is scanned again in the three fluorescent channels, giving the total protein signal (EGFP/A488 signal) and the relative amount of target bound (Cy5 signal). Each zinc finger fusion template was tagged with Cy3 during the final PCR step, and though signal from this channel was captured in each scan, it was not factored into downstream binding-specificity analyses, primarily because little Cy3 signal was detected as being bound by the ZF proteins and normalization for protein amount was performed with the EGFP fluorescence.

#### Image Analysis and Affinity Value Calculations

Images acquired from the experiment were processed using a 1024 unit detection array in GenePix v6.0. Raw tif files from the ArrayWoRx scanner were loaded into the GenePix software, and using the grid detection tool, an array of 1024 circular areas was snapped onto the EGFP spots detected in the A488 scan after washing. Small adjustments to the grid were performed by hand for poorly detected locations. Mean and median fluorescence measurements were taken in each fluorescent channel before and after washing. In addition, local background measurements were taken by dragging the detection array off the button valve locations into the space just outside the reaction chamber. For each scan, each circular area in the array was background corrected using its own local background measurement. Datapoints were filtered to ensure that EGFP levels were at least 500 AFU or higher and Cy5 target levels at least 1000 AFU or higher. These filtered, and background-corrected data points were used to calculate relative affinity values:

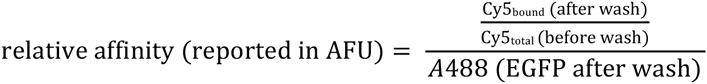

These normalized affinity values were used to compare ZF affinity to various targets across different experiments. In general, at least 2 data points were averaged together to arrive at a single affinity value (as in the heatmaps generated in Figure 1f and 2a,b).

#### Protein Modeling Methods

The Rosetta software suite^29^ was used to model the Zif268 (PDBid-1A1L^39^) alanine mutants in the presence of the DNA target consensus sequence. First, we computed differences in Rosetta energy units regarding the protein-DNA interaction^40^ caused by the mutations (ΔΔG_bind_= Rosetta Binding Energy_*mut*_ –Rosetta Binding Energy_*WT*_), the Rosetta Binding Energy for intermolecular interactions is computed by the difference of between the energy of the complex and the sum of the energies of the two molecules apart (Rosetta Binding Energy = Rosetta Energy_complex_ –(Rosetta Energy_partner1_ + Rosetta Energy_partner2_). To perform such calculations we used the Rosettascripts application^41^ with which mutants were generated and side chains in contact with DNA base pairs were subjected to energy mimization by sampling small deviations of the dihedral angles of the rotamers present in the crystal structure.

The stability calculations for Zif268 were also performed using Rosettascripts the differences in stability were computed through the difference in Rosetta energy between the mutants and the wild-type proteins (ΔΔG_stab_= Rosetta Energy_mut_ – Rosetta Energy_WT_). These simulations where performed in the presence of the zinc ions and all side chains were allowed to sample different rotamers and where energy minimized by sampling small deviations in the dihedral angles. For each mutant, 50 structural models were generated and the average of Rosetta energies is reported. Detailed command lines are presented below.

Rosetta command lines and scripts:

Rosetta version from Github repository -2662b747e67cf11cd76e6dedf2e9ff48cfefcd7c

Protein-DNA binding energy calculations

Command line:

~~~
rosetta_scripts.linuxgccrelease -s 1A1L_0001.pdb -parser:protocol prot- dna_script.xml -nstruct 3 -ignore_unrecognized_res
~~~

Rosettascripts code:

~~~
<ROSETTASCRIPTS>
  <TASKOPERATIONS>
    <InitializeFromCommandline name=IFC/>
    <IncludeCurrent name=IC/>
    <RestrictDesignToProteinDNAInterface name=DnaInt base_only=1 z_cutoff=3.0 dna_defs=B.1.GUA/>
    <OperateOnCertainResidues name=AUTOprot>
      <AddBehaviorRLT behavior=AUTO/>
      <ResidueHasProperty property=PROTEIN/>
    </OperateOnCertainResidues>
    <OperateOnCertainResidues name=ProtNoDesign>
      <RestrictToRepackingRLT/>
      <ResidueHasProperty property=PROTEIN/>
    </OperateOnCertainResidues>
    <OperateOnCertainResidues name=DnaNoPack>
      <PreventRepackingRLT/>
      <ResidueHasProperty property=DNA/>
    </OperateOnCertainResidues>
  </TASKOPERATIONS>
  <SCOREFXNS>
    <DNA weights=dna/>
  </SCOREFXNS>
  <FILTERS>
  </FILTERS>
  <MOVERS>
    <DnaInterfacePacker name=score scorefxn=DNA
task_operations=IFC,IC,AUTOprot,ProtNoDesign,DnaInt probe_specificity=1 binding=1/>
  </MOVERS>
  <PROTOCOLS>
    <Add mover_name=score/>
  </PROTOCOLS>
</ROSETTASCRIPTS>
~~~

Protein stability calculations Command line:

~~~
rosetta_scripts.linuxgccrelease -s 1A1L_noDNA.pdb -parser:protocol prot_stab.xml -nstruct 50 -ignore_unrecognized_res -ex1 -ex2 - extrachi_cutoff 5 -in:auto_setup_metals
~~~

Rosettascripts code:

~~~
<ROSETTASCRIPTS>
  <TASKOPERATIONS>
    <ReadResfile name=rrf filename=resfile/>
  </TASKOPERATIONS>
  <SCOREFXNS>
    <scorefxn1 weights=talaris2013>
    <Reweight scoretype=“atom_pair_constraint” weight=1.0/>
    <Reweight scoretype=“angle_constraint” weight=1.0 />
    </scorefxn1>
  </SCOREFXNS>
  <FILTERS>
  </FILTERS>
  <MOVERS>
    <PackRotamersMover name=packrot task_operations=rrf scorefxn= scorefxn1/>
    <Prepack name=ppk jump_number=0 scorefxn= scorefxn1/>
    <MinMover name=sc_bb_min bb=0 chi=1 scorefxn= scorefxn1/>
  </MOVERS>
  <PROTOCOLS>
    <Add mover_name=packrot />
    <Add mover_name=ppk />
    <Add mover_name=sc_bb_min />
  </PROTOCOLS>
</ROSETTASCRIPTS>
~~~

